# Modeling speech adaptation to altered sensory feedback through continuous learning of internal sensory predictions

**DOI:** 10.1101/2024.11.26.625353

**Authors:** Benjamin Elie, Juraj Šimko, Alice Turk

## Abstract

The paper presents a version of an optimization-based model of speech production that reproduces key acoustic and articulatory features of motorsensory adaptation to altered sensory feedback. In the presented approach, the mechanism of motorsensory adaptation is based on regular updates, based on the sensory feedback perceived by the speaker, of two of the speakers’ internal models used for computing (near)-optimal articulation. These internal models, modeled as separate Artificial Neural Networks, are 1) a model that predicts the acoustic consequences of motor (articulatory commands) and 2) a model that predicts the somatosensory sensations from given motor commands. The paper presents simulations of adaptation experiments that successfully reproduce key acoustic and articulatory features of motorsensory adaptation of speech to altered sensory feedback. These include gradual and incomplete motorsensory adaptation when the auditory (or the somatosensory) feedback is suddenly altered (F1-shifted for the altered auditory feedback, forced jaw movement for altered somatosensory feedback). The presented simulations also show that the rate and magnitude of adaptation behavior depend on a small number of parameters. Variation in the values of these parameters can potentially explain inter-speaker differences in terms of adaptation behavior, including sensory preference.

## Introduction

Turk & Shattuck-Hufnagel [1, 2] recently proposed an XT/3C (Phonology-Extrinsic-Timing-Based, Three-Component) approach to modeling speech production based on symbolic phonological representations and phonology-extrinsic timing. It has 3 processing Components: 1) Phonological Planning, 2) Phonetic Planning, and 3) Motor-Sensory Implementation. We have recently begun developing a computational implementation of the Phonetic Planning Component, which assumes an Optimal Control based process that finds the lowest cost movements that accomplish phonological goals (cf. [3–5]). We have shown that this approach can account for various aspects of speech, including durational and spatial correlates of prominence, including reduction and centralization of unstressed vowels and lenition of consonants [6], different levels of hypo and hyper-articulation [6–8], and articulatory and acoustic effects of Lombard speech [9].

In the current paper, our goal is to model sensorimotor compensation and adaptation within this framework. When exposed to perturbed auditory feedback, speakers often compensate on the next production by modifying their speech production in the direction opposite to the perturbations. Compensation is usually incomplete, and takes a number of trials before it stabilizes. Compensatory behavior has been experimentally observed when the feedback is formant-shifted [10–24], F0-shifted [25–31], or temporally altered [32]. After alterations to auditory feedback are turned off, speakers’ productions do not return to pre-perturbation patterns immediately, but instead do so gradually. Articulatory compensation has also been observed when somatosensory feedback is altered [13, 33–35]. These findings show that sensory feedback plays a role in speech production and speech articulatory planning.

This study will allow us to begin to address the third processing component of XT/3C [1, 2], namely the Motor-Sensory Implementation Component, and to further develop the Phonetic Planning component to account for adaptation behavior. The Motor-Sensory Implementation Component is envisaged as the stage when productions actually occur, and are monitored and adapted to ensure that as far as possible, the goals for the utterance are met. Compensatory behavior occurs in reaction to sensory effects of produced speech, and must involve mechanisms in the Motor-Sensory Implementation Component that monitor the (altered) feedback. Experience with this feedback in turn affects the plans for subsequent productions, namely plans that are made in the hypothesized Phonetic Planning component in XT/3C [1, 2].

We begin with a brief review of existing approaches to modeling compensation and adaptation behavior, which broadly speaking, can be classified as either correction-based, or based on adaptation of internal models of relationships among articulation, sensory consequences, and phonemic goals.

We then present a more detailed view of the compensation and adaptation behavior that we propose to model, followed by our adaptation-based account of this behavior. We provide simulations that show that XT/3C can provide an account using an architecture that is independently motivated by XT/3C’s assumption that speakers intend to signal acoustic cues to phonological categories, and by XT/3C’s optimization approach to Phonetic Planning. However, the internal models that we previously used to account for phonetic behavior in [6–9] must be supplemented with internal models that relate articulation to somatosensation (as e.g., in [36–38]), as well as somatosensation to phoneme recognition likelihood [36, 37].

Finally, we compare fits of our model predictions to experimental data with those of SimpleDIVA [39], which provides a correction-based approach. We show that results of the two approaches are very similar.

### Brief review of existing approaches

Several existing models are able to reproduce observed compensation and adaptation behavior. These models provide hypotheses about the mechanisms that speakers use in producing this behavior. Available models include SimpleDIVA [39], derived from the DIVA model [40–42], Bayesian GEPPETO [36, 37, 43], derived from the GEPPETO model [44], and FACTS [38, 45].

Both DIVA and FACTS provide *correction-based* mechanisms: they use auditory and somatosensory feedback to monitor and update feedforward commands, where the update is based on a comparison between the target sensory feedback (either auditory or somatosensory) and received sensory feedback. Feedforward commands are thus corrected (updated) when discrepancies are detected between expected sensory (somatosensory and auditory) targets and sensory feedback. The correction is proportional to the difference between the target (or predicted) sensory feedback and the actual sensory feedback received by the speaker. FACTS [38, 45] is similar to DIVA [40–42] in that it uses an error detection mechanism to update feedforward commands, but because it uses an internal model of the relationship between an efference copy of motor commands and their sensory consequences, it additionally updates the somatosensory and auditory prediction models when an error is detected. The correction gain is based on the prediction error.

Bayesian GEPPETO [36, 37, 43] provides a mechanism that is solely based on *adaptation*. Bayesian GEPPETO is a probabilistic generative model that generates feedforward motor commands by sampling from a distribution of motor commands associated with the auditory and somatosensory characterization of target phonemes. The probability distribution from which the feedforward motor commands are sampled is the product of several probability distributions, including the probability of auditory and somatosensory consequences given motor commands, the probability of acoustic and somatosoensory characteristics given the target phonemes, and matching constraint functions that are used to compare the sensory-motor predictions and the sensory characterization of phonemes. In order to model adaptation behavior using Bayesian GEPPETO, the authors in [36, 37] assumed that the probability distributions of the auditory and somatosensory consequences given a set of motor commands are updated based on discrepancies between predicted and perceived sensory feedback. As a consequence, this affects the probability distribution of feedforward motor commands associated with each phoneme, yielding new acoustic realizations of target phonemes. Here, adaptation occurs because motor commands chosen from the updated probability distribution are more likely to generate auditory sensations that correspond to the characterization of the target phoneme in the auditory space. This is because the internal motor command to auditory (or somatosensory) sensation has been updated and has learned the new (artificial) articulatory to sensory mapping the sensory perturbation.

### Modeling framework

In this paper, we present an adaptation-based account of compensation and adaptation behavior, that does not involve explicit detection of a discrepancy between predicted and actual sensory feedback to update feedforward commands and internal models of sensory predictions in order to correct for the discrepancy. Instead, our approach is a purely *adaptation-based* approach, involving continuous, permanent, and automatic learning and adaptation to an evolving environment, which is conceptually similar to the framework provided in the Bayesian GEPPETO papers [36, 37]. The main difference between our approach and GEPPETO is that the adaptation model we propose is based on Optimal Control Theory (OCT) [46, 47]: We assume that speakers plan articulatory movements that satisfy conflicting linguistic and extra-linguistic requirements [6–9], that is, that speakers minimize a cost function that includes weighted costs of effort and intelligibility^1^.

The version of our optimization approach presented here includes an internal forward model that predicts the somatosensory consequences of articulatory commands, in addition to the internal forward model predicting the acoustic consequences of articulatory commands included in the previous versions. We show that compensation and adaptation behavior can be accounted for through automatic updating of the assumed internal models of the relationship between articulation and sensory consequences, both acoustic and somatosensory. This type of internal model is independently motivated within our optimization approach; internal models which predict sensory consequences of movement are required because planned articulations are evaluated according to intelligibility costs related to the prediction of recognition of target phonemes which in turn are based on the prediction of acoustics and somatosensory sensations of the candidate motor commands. As such, for both the auditory and somatosensory branches, we assume a 2-part mapping: 1) between motor command and the corresponding sensory consequences (acoustics for the auditory branch, somatosensation for the somatosensory branch), and 2) between the corresponding sensory consequences and predicted probability of recognition of target phonemes. We assume these two types of internal models because they may be learned at different times of life, possibly at different rates, and possibly based on different information sources. For example, the models that relate motor commands to sensory consequences can start developing as soon as a baby begins to vocalize, and are based exclusively on the speaker’s own experience; on the other hand, the model of the relationship between acoustics and the probability of recognition might only start developing once the infant starts forming sound categories, and could potentially include information from other talkers (e.g. analogous to an exemplar-based, hybrid model of speech perception [51–53]).

The proposed overall cost function *J* to be minimized is as follows:

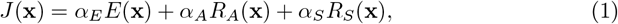

where *E, R*_*A*_ and *R*_*S*_ are the articulatory effort cost, and the predicted recognition costs associated with acoustics and somatosensory sensations, respectively. All are functions of the vector **x** that contains articulatory information, and the optimization task is to find the vector **x** that minimizes overall cost *J* (**x**).

The relative importance of the often mutually conflicting requirements enforced by these cost components is specified by the weights *α*_*E*_, *α*_*A*_, and *α*_*S*_, assigned to the least effort and the maximal auditory and somatosensory based intelligibility requirements, respectively. The recognition costs *R*_*A*_(**x**) and *R*_*S*_(**x**) in the cost function *J* (**x**) capturing a *non*-intelligibility of the utterance are computed using, firstly, internal forward models that predict the acoustic (for *R*_*A*_) and somatosensory (for *R*_*S*_) consequences of motor commands (included in **x**), and secondly, probabilistic models that associate a probability of recognition of the target phoneme given the predicted acoustic (for *R*_*A*_) and somatosensory (for *R*_*S*_) vectors.

In order to model sensorimotor adaptation of speech to altered auditory feedback, the optimization-based approach needs to change either the weights assigned to the different costs used in Eq. (1) and/or modify the models used to compute *E*(**x**), *R*_*A*_(**x**), and *R*_*S*_(**x**). This is because, without modification of Eq. (1) or its parameter values, and assuming that the speaker has no information about the new relationship between produced acoustic (or somatosensory) and altered sensory feedback during the adaptation phase in sensorimotor adaptation experiments, the solution that minimizes Eq. (1) would remain the same, that is, with no adaptation behavior. In this paper, we propose an approach which is based on automatic updates of the speaker’s internal forward models, namely those used to compute the cost function in Eq. (1), based on the speaker’s sensory feedback. In this approach, updating the model of the internal relationship between articulation and sensory characteristics would mean that candidate motor commands would have different Recognition costs; optimal articulations are therefore likely to change.

### Modeling objectives

The features of sensorimotor adaptation that we attempt to reproduce with our approach are detailed in this section along with possible accounts of these phenomena. These include gradual adaptation (Objective A), incomplete compensation (Objective B), interspeaker variability (Objective C), changes to unaltered formants during adaptation (Objective D), and adaptation to somatosensory perturbations (Objective E). Gradual adaptation and incomplete compensation are illustrated in Figure 1.

**Figure 1.**
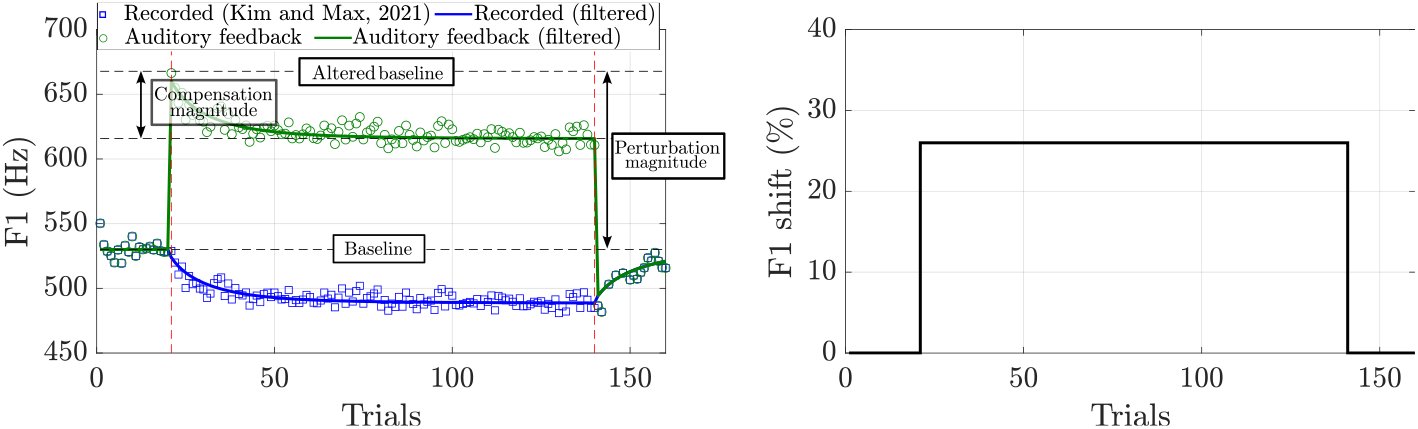
Example of experimental data that show adaptation to altered auditory feedback. Experimental data are taken from Kim and Max [21]. The experiments used proportional perturbations of F1, where the (upward) shift in % is shown on the right panel. The left panel shows the evolution of the recorded F1 (blue squares), and the auditory feedback received by the speaker (green circles), namely the first formant frequency F1 (altered during the perturbation phase). Solid lines represent a smoothed (filtered) version of the recorded and heard F1. The baseline corresponds to the mean value of the produced F1 before the perturbation phase (here, trials 11 to 20, with a baseline value of 530 Hz). The altered baseline corresponds to the value in Hz of the baseline when the perturbation is applied. The perturbation magnitude is the difference, in Hz, between the altered baseline and the baseline value. The compensation magnitude is the difference in Hz between the value of the auditory feedback received during the adaptation phase that is closest to the baseline, and the altered baseline.

#### Objective A: Gradual adaptation

The process of motor-sensory adaptation is generally not immediate. As shown in the left panel of Figure 1, which presents experimental data from [21], once sensory feedback is altered it takes several trials for subjects to reach their maximal adaptation response. The maximal adaptation response is the production that generates altered sensory feedback closest to the sensory feedback produced before sensory perturbation was applied. In Figure 1, maximal adaptation occurs around trial 70. For speech, gradual adaptation is observed, for instance, in auditory perturbation experiments in [12, 14, 16, 21, 24, 54]. The left panel of Figure 1 shows that it took around 10 trials (10 target word productions) after the beginning of the auditory perturbation phase for speakers to produce an adaptation response close to the maximal adaptation response.

This gradual adaptation in auditory perturbation based experiments has been reproduced by FACTS [45] and SimpleDIVA [39]. In SimpleDIVA, the speed at which speakers tend toward their maximal adaptation response is directly included in the model and specified by the learning rate parameter. This learning rate parameter corresponds to the proportion, assumed to be constant throughout the experiments, of the feedback-based corrective command to be added to the new feedforward command. In FACTS [45], realistic adaptation responses were achieved by using adaptive Kalman filters to control and monitor the gain to apply to the task state correction, i.e. the correction applied to the estimate of the current state of the articulators, when sufficiently large discrepancies (higher than a specific threshold) between the predicted and the actual received sensory feedback were detected. Similarly to the role of the learning rate parameter in SimpleDIVA [39], the adaptive Kalman filter gains impacts the adaptation rate: faster adaptation rates can be achieved in FACTS with larger adaptive Kalman filter gains, simply because more correction is applied to the updated task state. Note that unlike SimpleDIVA, the modified version of FACTS in [45] allows feedback-based correction gains to vary during the experiments: in FACTS, the gain can be increased when the discrepancy between predicted and received sensory feedback is higher than a pre-defined threshold. This increase in feedback-based correction gains allows larger and faster update in case of large and obvious detected prediction “error”.

In this paper, we propose an approach that does not require a comparison between predicted and actual sensory feedback. Our approach uses an automatic and permanent update of the speaker’s internal models that predict the acoustic and articulatory consequences of motor commands. Our assumption is that building these internal models is part experience with vocalization, including speech: these internal models are continuously trained and updated during the speaker’s lifetime based on production and sensory feedback, regardless of whether the model predictions are accurate. Consequently, during adaptation to altered auditory feedback, the motor command to acoustics internal model learns the new (artificial) relationship between motor commands and the acoustic consequences. We hypothesize that the completion of the learning process may take several trials to complete, resulting in gradual adaptation. Similarly, when the feedback perturbation is removed, we expect our approach to account for a gradual return to the baseline.

#### Objective B: Incomplete compensation

Experimental studies reported that compensation to feedback perturbations is usually incomplete (see e.g. [12, 19, 21, 23, 54]). Kitchen *et al*. [54] reported compensation magnitudes that range from 20-40% of the auditory perturbation. In the particular example presented in the left panel of Figure 1, using data from [21], the compensation ratio (defined as the ratio between compensation magnitude and perturbation magnitude) is 37.7%.

In FACTS, as in [45], incomplete compensation is explained by convergence between actual and predicted auditory output. More precisely, the (altered) auditory feedback and the predicted auditory output converge during the adaptation phase because 1) the altered auditory feedback goes towards the baseline because of the speaker’s compensation and 2) internal models used to predict the auditory output from motor commands are updated so that new predictions go in the same direction as the perturbation, leading to an eventual convergence. For instance, for a raised-F1 type of perturbation, new predictions from the updated internal model will estimate higher F1 than before perturbation, because the earlier predictions were evaluated as erroneously too low. Subsequently, the feedforward command will be updated in order to produce lower F1 to compensate for the unexpectedly high F1 perceived under altered auditory feedback. As a result, the F1 perceived altered auditory feedback will be lower on a subsequent trial. This gradually increasing predicted F1 and the gradually decreasing perceived F1 eventually converge and meet somewhere in the F1 space between the baseline and the altered baseline. This convergence nullifies the discrepancies between predicted and actual auditory feedback before full compensation is reached, removing the need for further adaptation by the speaker and yielding incomplete compensation.

Since our approach does not use a direct comparison between sensory prediction and received sensory feedback, this convergence mechanism does not apply. Instead, we rely on updates of internal models that relate motor commands to sensory consequences. This approach requires models that relate both 1) motor commands to acoustics, and 2) motor commands to somatosensation. Because our previous model had a predicted recognition cost that determined the probability of phoneme recognition based only on its acoustics consequences, as in [6–9], there would be no limit to compensation, as the auditory internal model would learn the artificially modified articulatory to acoustic mapping and eventually find the solution that minimizes the cost function with a predicted acoustic output that corresponds to the baseline. This purely auditory-based kind of recognition optimization is analogous to the purely auditory planning process in Bayesian GEPPETO [37], which also predicts full compensation because of the absence of any constraint on the somatosensory characterization of target phonemes. In order to obtain incomplete adaptation, we propose to include somatosensory information in the cost function. In doing so, we predict that the solution that minimizes the cost function will be a trade-off between full acoustic compensation (satisfying the auditory objective) and remaining close to the somatosensory sensations associated to the target phonemes (satisfying the somatosensory objective). This solution is inspired by Bayesian GEPPETO [37], in which incomplete compensation is achieved using an extension of the purely auditory planning process, namely the fusion planning process, which uses both the auditory and somatosensory characterization of target phonemes. That is, incomplete adaptation is explained by a balance of somatosensory and auditory adaptation: the articulatory solution for complete adaptation would generate somatosensory feedback too far from the expected one. In this paper, we propose to add an extra cost in the cost function that accounts for the speaker’s somatosensory characterization of phonemes, as in Eq. (1).

#### Objective C: Inter-speaker variability

Sensorimotor adaptation abilities have been shown to be strongly speaker dependent, even among neurotypical speakers with similar cognitive characteristics and language backgrounds [23, 24, 54]. Speakers also tend to compensate more or less depending on the nature of the sensory perturbation. This has been called *sensory preference*, reported in [13]: Some speakers compensate more for the auditory perturbations while others compensate more for somatosensory perturbations. In addition to these inter-speaker sources of variability, many studies have found that adaptation behaviors can differ across groups of speakers, based on age [24], auditory acuity [19, 23], language (L1 or L2 speakers) [12, 23], cognitive ability [23], speech disorders (e.g. stuttering [21] and speech sound disorder [24]), or Parkinson’s disease [14].

Two variants of Bayesian GEPPETO have been proposed in [37] to model sensory preference. One approach specifies sensory preference by modulating the relative precision of the characterization of speech motor goals in both the auditory and the somatosensory spaces, namely by narrowing or widening the target regions of phonemes in their respective sensory spaces. More precisely, if the target regions are, let say, much narrower in the auditory space, compared to the target regions in the somatosensory space, perturbations in the auditory space will result in larger involvement of the auditory pathway in the planning process, and, conversely, less involvement of the somatosensory pathway. This is because large distributions (such as in the somatosensory space, in this case) allow for greater deviation from the center of the distribution. The other approach specifies sensory preference by modulating the matching constraints of the auditory and somatosensory pathways when comparing motor-sensory predictions and sensory characterization of phonemes. More precisely, the modulation of these matching constraints as for effect to allow more or less deviation from the center of the distribution of the target phoneme in the considered sensory space. For instance, if the matching constraints allows larger deviations in somatosensory space than in auditory space, it is expected than the speaker will compensate more for auditory perturbations than for somatosensory perturbations. Both approaches lead to very similar results.

Since SimpleDIVA has been designed to fit experimental data (and their variability), it naturally accounts for inter-speaker variability. Its use on experimental data showed that inter-speaker variability is explained in the SimpleDIVA framework by variability in feedback-based correction gains applied to the auditory and the somatosensory subsystems [24, 39].

As shown in our previous papers [6–9], modifying the weights of the cost function, as in Eq. (1), results in different characteristics of the output speech. A first source of inter-speaker variability in terms of adaptation behavior may then stem from different weights encoded in each speaker’s speech system. In addition, we hypothesize that characteristics of the internal sensory prediction models are speaker-dependent and explain the observed inter-speaker variability. These potentially different characteristics include the architecture and sizes of the Artificial Neural Networks that define the internal models, the amount of training data (exposition to language), accuracy of the perception of sensory feedback (e.g., poor auditory or somatosensory acuity lead to poor performance of the internal model because of inaccurate training data), as well as the model’s learning rate.

#### Objective D: Changes in unaltered formants during adaptation

Most experimental studies about adaptation to formant-shifted auditory feedback only analyzed the modification of the production of the altered formant or formants, and ignored the behavior of unaltered formants. That is, most studies have reported variations across trials of only F1 for F1-shifted auditory feedback. The small number of studies which have also measured the behavior of other formants have reported that these other formants were also impacted during adaptation [11, 14, 16, 29]. Basically, F1-raised auditory feedback induced both F1 lowering and F2 raising. However, changes in unaltered formants are not always observed: no systematic change in F2 was observed in [55] for F1-raised auditory feedback among 18 participants that adapted to altered F1 auditory feedback.

These observations cannot be explained by purely acoustic target based models if based on single formant targets, such as in SimpleDIVA [39] or the forward/inverse model in [56]. This is because these models assume that the formant targets which are not altered remain the same during the adaptation process. As a result, they predict that planned articulatory commands will compensate for formants which have been altered, but not for other, unaltered, formants.

This aspect is not explicitly addressed in the studies using Bayesian GEPPETO, such as in [36, 37]. However, the results of the simulations presented in [37] suggest that, in the presence of F1-shifted auditory feedback, the predicted acoustic output of the fusion model exhibits a change in both F1 and F2. However, the authors do not discuss this question. FACTS [45] is able to successfully reproduce similar changes in F1 and F2 to the adaptation behavior experimentally observed in [14]. When discussing these results, the authors provide possible explanations for mechanisms that could affect non-altered formants during adaptation. One possible explanation discussed by the authors is that targets, in FACTS [38], are defined in the vocal tract constriction location and degree task space (namely as tract variables defined by the gestural score). Updating the articulatory constriction tasks during adaptation yield simultaneous changes in all formants because modifying articulatory configurations affects all formants simultaneously. In addition, there are biomechanical constraints on articulatory configurations that could prevent speakers from modifying a single formant independently of the others.

Our approach uses symbolic phonological representations as goals: Articulatory movements are planned to satisfy conflicting tasks of phoneme recognition and low articulatory effort. Formant-shift based perturbation of auditory feedback will result in modification of the probability of recognition of the target phonemes (as internalized by the speaker). As such, formants are not fully specified as targets: we expect compensation involves finding new articulatory configurations that generate target phonemes (as heard by the speaker) which are as likely to be recognized as before the perturbation and which still satisfy the trade-off between intelligibility and low articulatory effort. Consequently, there is no hard constraint on the unaltered formant frequencies and these can potentially be modified during the compensation process.

#### Objective E: Adaptation to somatosensory perturbations

One objective of this paper is also to show that our model is also able to simulate the kind of sensimotor adaptation observed in [13] using the same architecture as for the adaptation to altered auditory feedback. Since our approach assumes symmetric auditory and somatosensory subsystems, simulation of adaptation to either (or both) auditory and somatosensory feedback perturbations should be very similar.

### An overview of our optimization-based model of speech production

The proposed approach is based on the assumption of a phonetic planning process that derives articulatory patterns that balance requirements of minimizing movement costs (articulatory effort in this paper, but time could also be included) while maximizing intelligibility of the produced utterance. During the optimization process, intelligibility is estimated using internal predictions of both acoustic and somatosensory consequences of a multitude of candidate, not yet realized articulatory actions. The presence of internal models that estimate recognition likelihood by a listener is thus a necessary consequence of the assumption of an online optimization process that forms the basis of the Phonetic Planning component in XT/3C.

The acoustic and somatosensory consequences of articulatory actions are subsequently used to predict probabilities of recognition of the planned phonemes by a listener. The mapping between sensory characteristics and probability of recognition is assumed to be fixed in the present version. In our current approach, the internal models mapping articulation to acoustic and somatosensory sensations are updated after each produced utterance based on the actual percepts of the speaker. This is similar to the approach presented in FACTS [45], where the definitions of task dynamics are updated when there is a discrepancy between predicted and actual output. However, unlike FACTS, we do not assume any specific error detection mechanism. Instead, the prediction models relating motor commands to sensation are updated after each production. Any alteration of perceptual (acoustic or somatosensory) feedback will thus result in a change of the relevant model, leading to a different output of the optimization process in subsequent steps. When, for example, speech output is modified by an experimenter by artificially shifting a formant upwards, the speaker will gradually internalize this change, and will compensate for the modification in subsequent utterances. So, unlike SimpleDIVA and FACTS models that account for adaptation phenomena through explicit modification of actual feedforward commands, our approach uses updates of internal models that are used to derive the optimal articulatory realizations.

Like GEPPETO, we define two internal forward models that map motor commands onto sensory spaces, namely an acoustic (auditory) model and a somatosensory model. The acoustic model, denoted *𝒜*, is used to predict the acoustic consequences of motor commands. The somatosensory model, denoted *𝒮*, is used to predict the somatosensory sensations that result from motor commands. This yields the following relationships:

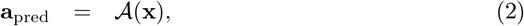

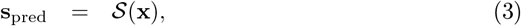

where **a**_pred_ and **s**_pred_ are the predicted acoustic and somatosensory vectors, respectively, given the vector of articulatory commands **x**. In our approach, the predicted sensory vectors **a**_pred_ and **s**_pred_ are subsequently used to estimate the probability of recognition of target phonemes.

For the sake of simplicity, we assume in this paper that the acoustic consequences are formant frequencies and that the somatosensory sensations are the tract variables. Discussing the choice of acoustic and somatosensory representations is left for future work. In this paper, both *𝒜* and *𝒮* are implemented in the form of MultiLayer Perceptrons (MLPs), namely feedforward fully connected Artificial Neural Networks (ANNs). Updating these internal models after each trial consists of updating weights of the corresponding MLP via fine-tuning through a few epochs using a motor command and sensory output pair which resulted from the previous trial. For instance, the input/output pair used to update *𝒜*^(*n*)^ after trial *n* consists of the vector of motor command **x**^(*n*)^ provided by the optimization process at trial *n* and the vector of formants 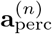 perceived by the (virtual) speaker at trial *n* (*i*.*e*., its auditory feedback, which is potentially perturbed during the simulated adaptation experiment). Similarly, the input/output pair used to update the somatosensory model *𝒮*^(*n*)^ after trial *n* consists of **x**^(*n*)^ and the somatosensory vector 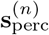 perceived by the speaker at trial *n*.

To generate the acoustic and somatosensory representations, *i*.*e*. the formant frequency and tract variables respectively, we use the Maeda model [57] in this paper. The Maeda model generates midsagittal shapes of the vocal tract using seven independent articulatory parameters, corresponding to the principal components that explain most of the observed variance in articulatory data. These are expressed in terms of standard deviations above or below the mean value, where the mean value (i.e. 0) corresponds to the parameter’s value in a neutral vocal tract position. The tract variables are computed following the technique detailed in [58], to which we add vocal tract length (VTL), computed as the length of the vocal tract midline from the larynx to the end of the lips. We used the Maedeep python library [59] implementation, which is a python version of the original VTCalcs module, written in C [60] to compute the formant frequencies associated with the vector **x** of Maeda parameters. For the sake of simplicity, we used the Maeda parameters (*i*.*e*., the seven independent articulatory parameters introduced above) as our vector of motor commands. Consequently, *𝒜* learns the mapping between Maeda parameter values and associated formant frequencies, while *𝒮* learns the mapping between Maeda parameter values and tract variable values.

The architecture of our model is represented in Figure 2. At each trial *n*, the optimization block plans a vector of motor commands **x**_*n*_ that minimizes the cost function, using the internal models *𝒜*^(*n−*1)^ and *𝒮*^(*n−*1)^, for a given target phoneme *p*. The optimization block uses the vector of motor commands of the previous trial **x**_*n−*1_ as the initial solution. The plant (here the Maeda model) is used to compute the produced acoustic vector 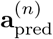 and the produced somatosensory vector 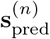. Depending on the nature and on the amount of the sensory perturbation, these vectors are modified (or not if no perturbation is applied) to generate the sensory feedback vectors 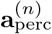 and 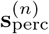. These sensory feedback vectors are used to update the weights of the internal models *𝒜*^(*n*)^ and *𝒮*^(*n*)^, which are then used to compute the solution of the optimization block at the next trial *n* + 1.

**Figure 2.**
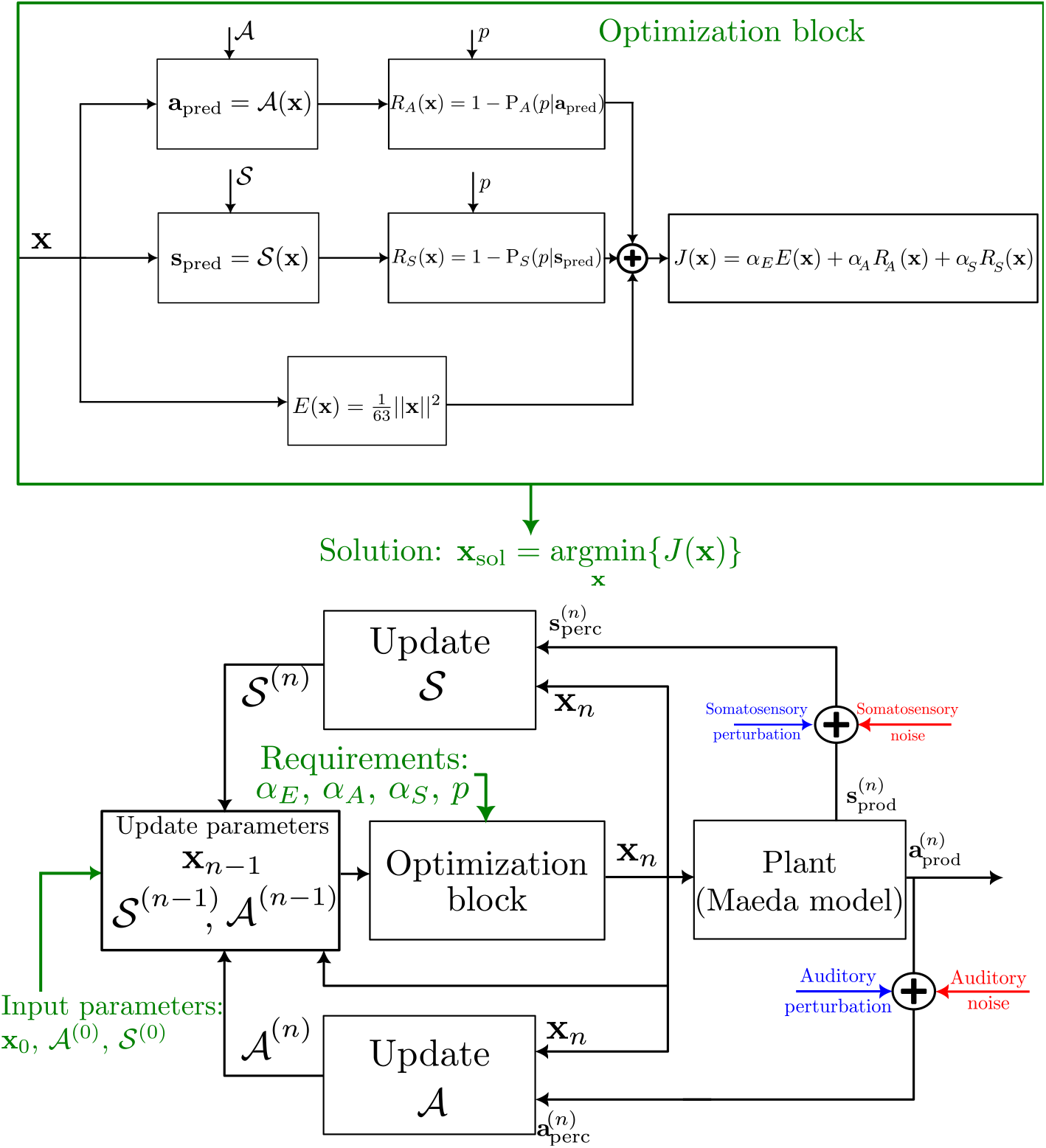
The architecture of our adaptation model. The top panel represents the optimization block. The bottom plot shows the general architecture of the model.

### The cost function

As introduced in Section Modeling framework, following [7–9], we assume a cost function that accounts for minimal articulatory effort and maximal intelligibility. The minimal model used in this paper generates static configurations. As a consequence, intelligibility is based on single frames of produced speech^2^. In the earlier versions of our OCT-based model [7–9], the intelligibility cost component was based on purely acoustic (auditory) consequences of articulatory movements. That is, the intelligibility cost component was assumed to be the speaker’s estimate of how intelligible a phoneme would be to a listener, given the acoustics only. As discussed in the introduction, we propose a new cost function that accounts for both the somatosensory and the auditory characterizations of phonemes. This translates into two independent costs related to an estimation of the probability that a phoneme will be recognized by a listener, namely one related to intelligibility given the acoustics, and one related to intelligibility given the somatosensory characterization of the target phoneme, as in Eq. (1). This is based on evidence that the speaker’s plan for articulatory movement is also based on their somatosensory characterization of phonemes. For instance, the phenomenon of *covert contrast* [62, 63] suggests that speakers with phonological disorder may use two distinct articulatory targets to distinguish a pair of phonemes (for instance alveolar and velar stop consonants) without making them acoustically distinguishable: their articulatory plan to distinguish these pairs of phonemes is predominantly based on distinct somatosensory output, as opposed to distinct acoustic consequences. The somatosensory characterization of phonemes in articulatory planning is taken into account in some models of speech production by two feedback loops, corresponding to auditory and somatosensory sensations, which are used to update feedforward commands, such as in DIVA [39–42], or in FACTS [38, 45]. In Bayesian GEPPETO [36, 37, 43], auditory and somatosensory characterizations of phonemes are modeled by a dual branch-based system of motor planning, including a subsystem that accounts for auditory goals, and another subsystem that accounts for somatosensory goals.

Still following [7–9], we assume that the cost of articulatory effort is related to the distance of articulators from their neutral position. Using Maeda’s articulatory model [57], this corresponds to computing the *𝓁*_2_*–*norm of the vector **x** of Maeda parameters:

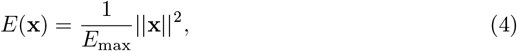

where *E*_max_ = 63 is used as a normalization factor to yield an effort cost with values ranging between 0 and 1 (63 being the maximal value of *E*(**x**), obtained when all elements in **x** are either -3 or +3). This normalization ensures that effort costs are in the same range of values as the recognition costs, whose values range from 0 to 1.

### Probabilistic models for computing intelligibility scores

The (predicted) recognition score *R* is the cost of not being intelligible, where intelligibility is modeled as the recognition probability of the target phoneme *p* given the articulatory vector **x**, hence:

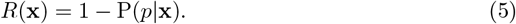

We consider two probabilistic models, denoted P_*A*_ and P_*S*_, which return the probability of the target phoneme given the predicted auditory and somatosensory sensations, respectively. These two probabilistic models are used to compute two distinct recognition costs *R*_*A*_ and *R*_*S*_, corresponding to the cost of recognition in the auditory and the somatosensory space, respectively, namely:

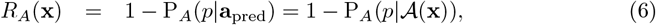

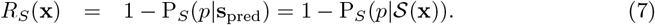

### Simulations

This section presents the simulations performed for this study, as well as fits to real experimental data.

### Training the internal forward models

In this paper, we used pre-trained internal forward models, denoted *𝒜*^(0)^ for the acoustic model and *𝒮*^(0)^ for the somatosensory model. They were trained using random generation of articulatory vectors **x** and with computation of associated formant frequency vectors **f** (for the acoustic model) and tract variable vectors **s** (for the somatosensory model) using the Maeda model [57]. We generated *N* articulatory vectors following a uniform random distribution between -3 and +3. We kept 90% for training and 10% for validation. Note that the associated formant vector of closed configurations (closed vocal tract) was set to the null vector (all formants at 0). We hypothesize that changing the architecture of internal models and/or the training data can modify the model’s flexibility to adapt and extrapolate to new observed data (auditory feedback), especially when these new observed data are far from what would be expected. That is, we believe that the size of the model (*i*.*e*., the number and size of the hidden layers) has a direct influence on the ability of the speaker to compensate for altered auditory feedback. Similarly, the amount of data used for training the internal models, corresponding to the past history of the speaker, can potentially affect the characteristics of the adaptation response, such as the maximal compensation or the number of trials that are necessary to reach that maximal compensation. In order to investigate the potential effect of the internal models’ characteristics on the adaptation behavior, we trained, for both internal models, various MLP networks with different number *M* of hidden layers of different sizes, and with a tanh activation function, during a maximal number of 100 epochs. Training was stopped when the validation loss was not improved for 5 successive epochs. For the simulations, we define nominal internal models as MLPs having the characteristics detailed in Table 1. Unless specifically mentioned, simulations are performed using these nominal internal forward models.

**Table 1.**
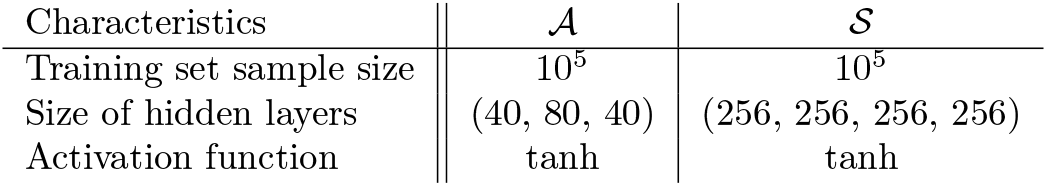
Architecture of the MLPs used for the nominal internal forward models.

### Training the internal probabilistic models

Similarly to the probabilistic model presented in [9], we used a Quadratic Discriminant Analysis (QDA) model trained on real formant values to predict the probability of vowels given a vector **a** containing the first 4 formant frequencies. In this paper, we trained the QDA model on formants of American-English vowels extracted from the Vocal Tract Resonance (VTR) database [64]. We did the same to train the somatosensory-vector-to-probability model *P*_*S*_, except for the input data being the vectors of tract variables **s**.

### Experimental data

In this paper, our simulations are based on participant data from the experimental study by Kim and Max [21]^3^. This is because we also aim to compare the results of our simulations with real experimental data from adaptation-based experiments. In this experimental study [21], speakers were asked to produce the American-English phoneme /ε/ in one of either *bed* or *pet* in sequences of 160 trials. During the first 20 trials, auditory feedback was not altered. For the next 120 trials, auditory feedback of speakers’ productions was altered by applying an upward shift of 400 cents (around 26%) to F1. Then, the alteration was removed for the last 20 trials. Our simulations are designed to simulate this experiment, namely producing the American-English phoneme /ε/ in sequences of 160 trials, with a perturbation function applied to F1 which is the same as the one used in [21]. This perturbation function is shown in the right panel of Figure 1.

## Methods

The presented simulations were conducted to investigate the influence of different aspects of our model on the adaptation process. For that purpose, we simulated several altered feedback experiments. These simulated experiments consisted of simulating the production of the English vowel /ε/ for 160 trials with a first formant frequency F1 altered as shown in Figure 1. Several parameters were modified for each simulation in order to investigate the effect of the weights of the cost function, as defined in Eq. (1), resulting in different degrees of hypo-/hyper-articulation, and also the effect of various characteristics of the pre-trained internal models, such as the learning rate, the number of training samples, the number, and sizes of hidden layers. Note that for optimization-based problems, the number of required weights is one unit less than the number of costs. This is because multiplying all weights by the same factor does not change the minimal solution. Consequently, only two weights are required. In this paper, we chose to normalize the auditory and somatosensory weights *α*_*A*_ and *α*_*S*_ such that they always sum as 1, namely *α*_*A*_ + *α*_*S*_ = 1.

Each simulation was performed as follows. We first chose specific pre-trained internal forward models, denoted *𝒜*^(0)^ and *𝒮*^(0)^, with specific learning rates 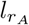 and 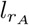 and a number of epochs *e*_*A*_ = *e*_*S*_ = 10 during updates, and fixed weights *α*_*E*_, *α*_*A*_, and *α*_*S*_ = 1 *− α*_*A*_. We used the Nelder-Mead algorithm [65] to run the initial optimization and estimate the optimal articulatory vector **x**^(0)^ that minimizes *J* in Eq. (1). For that first iteration, we used the pre-trained internal models *𝒜*^(0)^ and *𝒮*^(0)^ to compute both intelligibility scores based on auditory and somatosensory sensations. The estimated articulatory vector **x**^(0)^ was used to compute the actually produced formant vector 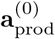 and produced somatosensory vectors 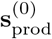. If there is no altered sensory feedback, the formant vector perceived by the speaker, denoted 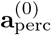, is simply 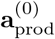. Similarly, the somatosensory vector perceived by the speaker, denoted 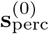, is simply the produced somatosensory vector 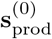. In that case, both **x**^(0)^ and 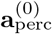 are used to fine-tune the internal model *𝒜*^(0)^ with *e*_*A*_ epochs run over this single sample. The fine-tuned version of the model is then *𝒜*^(1)^, and is used to estimate the new optimal articulatory vector **x**^(1)^, similarly to **x**^(0)^, and so forth until end of the simulations. Updating *𝒮*^(0)^ is done in similar fashion, except that the input/output pair used for fine-tuning is **x**^(0)^ and 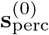. At the end of each iteration *n*, the internal models *𝒜*^(*n*)^and *𝒮*^(*n*)^ are obtained by updating the weight of *𝒜*^(*n−*1)^ and *𝒮*^(*n−*1)^ over *e*_*A*_ and *e*_*S*_ epochs, respectively, using only the samples **x**^(*n*)^ and 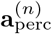 (respectively 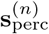). However, in the case of altered auditory feedback, the perceived formant vector at iteration *n* 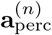 is an altered version of the actually produced formant vector 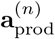, as:

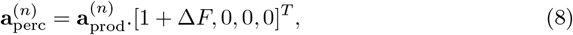

where Δ*F* as a function of the trial *n* is shown in Figure 1. That is, the alteration concerns only the first formant, the other formants being perceived as they are actually produced by the speaker.

Additionally, we compare the results of our simulations with those of the SimpleDIVA model. We chose SimpleDIVA as the baseline model for comparison because this model has been designed to fit experimental data. For that purpose, we fit our model and a 2-parameter version of SimpleDIVA to the real experimental data presented in [21]. That is, for a given speaker, we found the pair of parameters 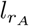 and *α*_*A*_ for which our model returned a F1 trajectory that minimized the RMSE with respect to the recorded F1 trajectory.

## Results of simulations with altered auditory feedback

This section presents the results of our simulations when the perturbation is applied to auditory feedback. These results show the influence of different parameters of our model, as well as the influence of the characteristics of the internal forward models, on the adaptation behavior.

### Illustration of a typical simulation

Figure 3 shows a typical example of the adaptation behavior returned by our simulations, with perturbations applied only to auditory feedback. The acoustic output visualized in Figure 3 (A) reproduces typical adaptation behavior to altered auditory feedback with perceived F1 shifted upwards between trials 20 and 140. The produced F1 (the blue solid line in Figure 3 (A)) moves downwards in the direction opposite to the perturbation. As observed experimentally in previous studies [12, 14, 16, 21, 24, 54], the adaptation is gradual, as it takes several trials to reach an altered auditory feedback (F1 value) close to the maximal compensation (defined as the lowest altered F1 value). Additionally, the compensation is not complete, as the minimal heard F1 (auditory feedback, in red) during the perturbation phase is higher than the baseline (gray dotted line), i.e., the F1 perceived before the perturbation was applied. This type of incomplete adaptation is also a feature commonly observed experimentally in adaptation studies (e.g., in [12, 19, 21, 23, 54]).

**Figure 3.**
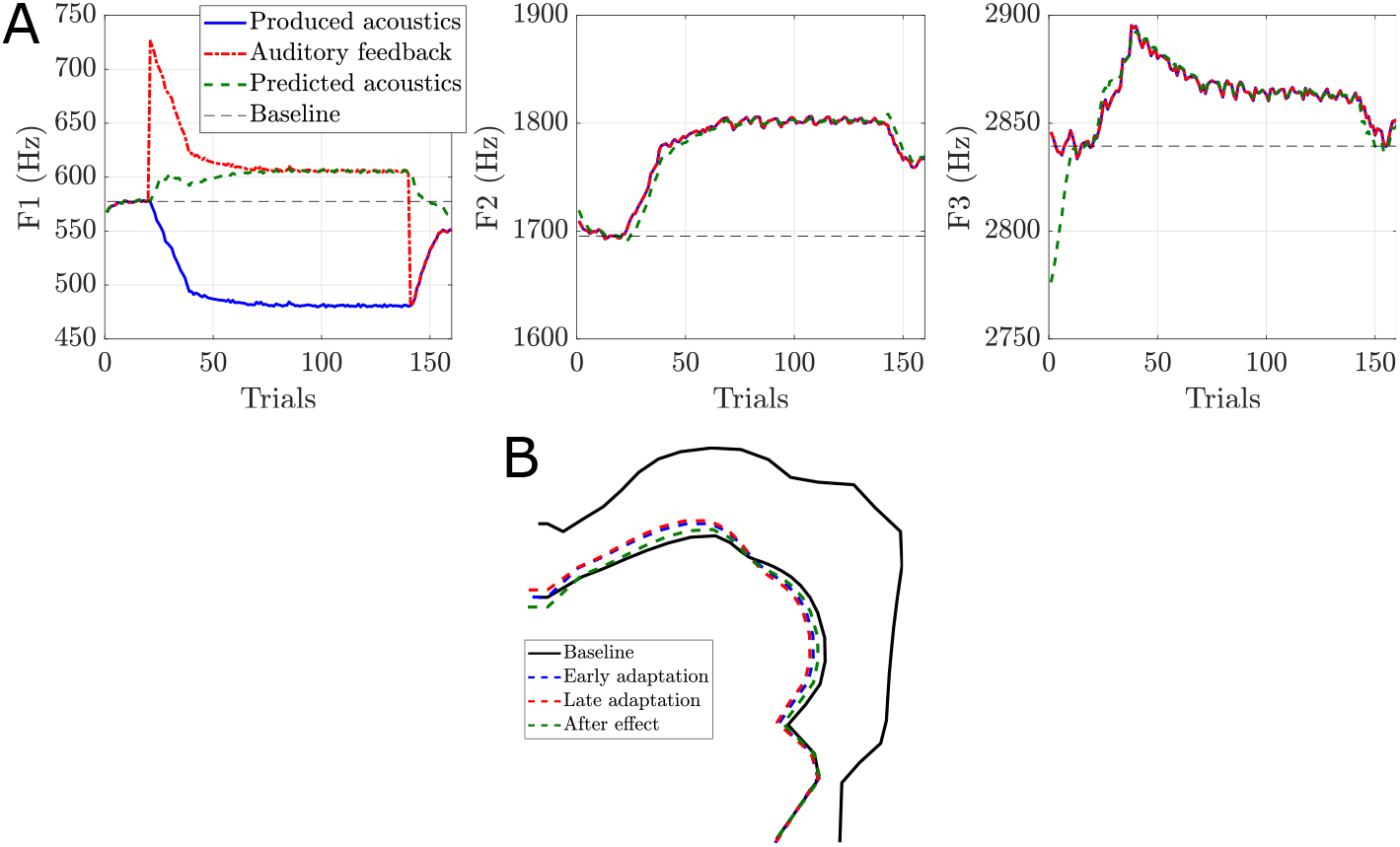
A typical example of simulated adaptation to altered auditory feedback. A) shows simulated acoustic output in separate panels for formant frequencies F1, F2, and F3, including produced acoustics (blue solid line), auditory feedback (dash-dotted red line), and acoustics predicted by the auditory internal model (dashed green line). The bottom plot B) shows the mean simulated shape of the vocal tract as returned by the Maeda model, averaged across the baseline phase (without perturbation, trials 11 to 20), the early adaptation phase (trials 21 to 80), the late adaptation phase (trials 81 to 140), and the end phase (after-effects, trials 151 to 160).

Figure 3 (A) also shows the acoustic output as predicted by the auditory internal model for each subsequent trial, based on the performed articulatory action. Within our optimization-based approach, it is this predicted acoustic output–rather than the actual “heard” realization–that is evaluated as the optimal realization of the speaker’s intention, i.e., that minimizes the cost function *J* in Eq. (1). In this example, the simulations updated the internal model predicting the acoustic consequence very early in the adaptation phase, around trial 25. While the predicted value of the acoustic consequence of the articulation largely stabilizes after this trial, these relatively constant predictions correspond to different, downward shifting actual productions. The internal model keeps being updated and drives behavioral adaptation until the gap between predicted and perceived acoustic consequences (dash-dotted red and dashed green lines) disappears at around trial 60. This gradual adaptation of the internal model of articulatory consequences of articulatory action thus gives rise to gradual behavioral adaptation.

Our simulations also predict that, in addition to F1, formants F2 and F3 are also modified during the adaptation phase; F2 and F3 both shift upward. The maximal upward shift of F2 is around 5% of the baseline value, which is in agreement with the F2 shift observed in [14]. During the adaptation phase, the simulated example in Figure 3 predicts that the speaker would raise their tongue and move it frontward, making the original /ε/ higher and more front, namely tending towards /e/ and /i/ (see Figure 3 (B)). This articulatory shift would result in lower F1 (because of adaptation to the altered F1), but would also increase F2.

This typical example of simulation illustrates that some of our modeling objectives presented in the introduction are fulfilled. This is because it shows that our approach is able to reproduce gradual (Objective A) and incomplete (Objective B) adaptation to altered auditory feedback by moving the altered formants in the direction opposite to the perturbation. In addition, the adaptation also modified other formants, namely F2 and F3, although they were not modified in the auditory feedback (Objective D).

### The influence of the somatosensory intelligibility preference on compensation

Figure 4 shows the results of our simulations for the American-English vowel /ε/ under formant-shifted perturbations shown in Figure 1, for different values of *α*_*S*_, the relative weight assigned to the somatosensory intelligibility. We remind the reader that the weight assigned to acoustic intelligibility is *α*_*A*_ = 1 *− α*_*S*_. This means that when *α*_*S*_ = 0 (left plot in Figure 4), only acoustic intelligibility is optimized, while when *α*_*S*_ = 1 (right plot in Figure 4), only somatosensory intelligibility is optimized, as *α*_*A*_ = 0. In these experiments, the weight assigned to the effort cost was 0. This was done to make sure that the observed effect was due to the different values of *α*_*A*_ and *α*_*S*_, and not due to a different ratio between one of those weights and the effort weight *α*_*E*_.

**Figure 4.**
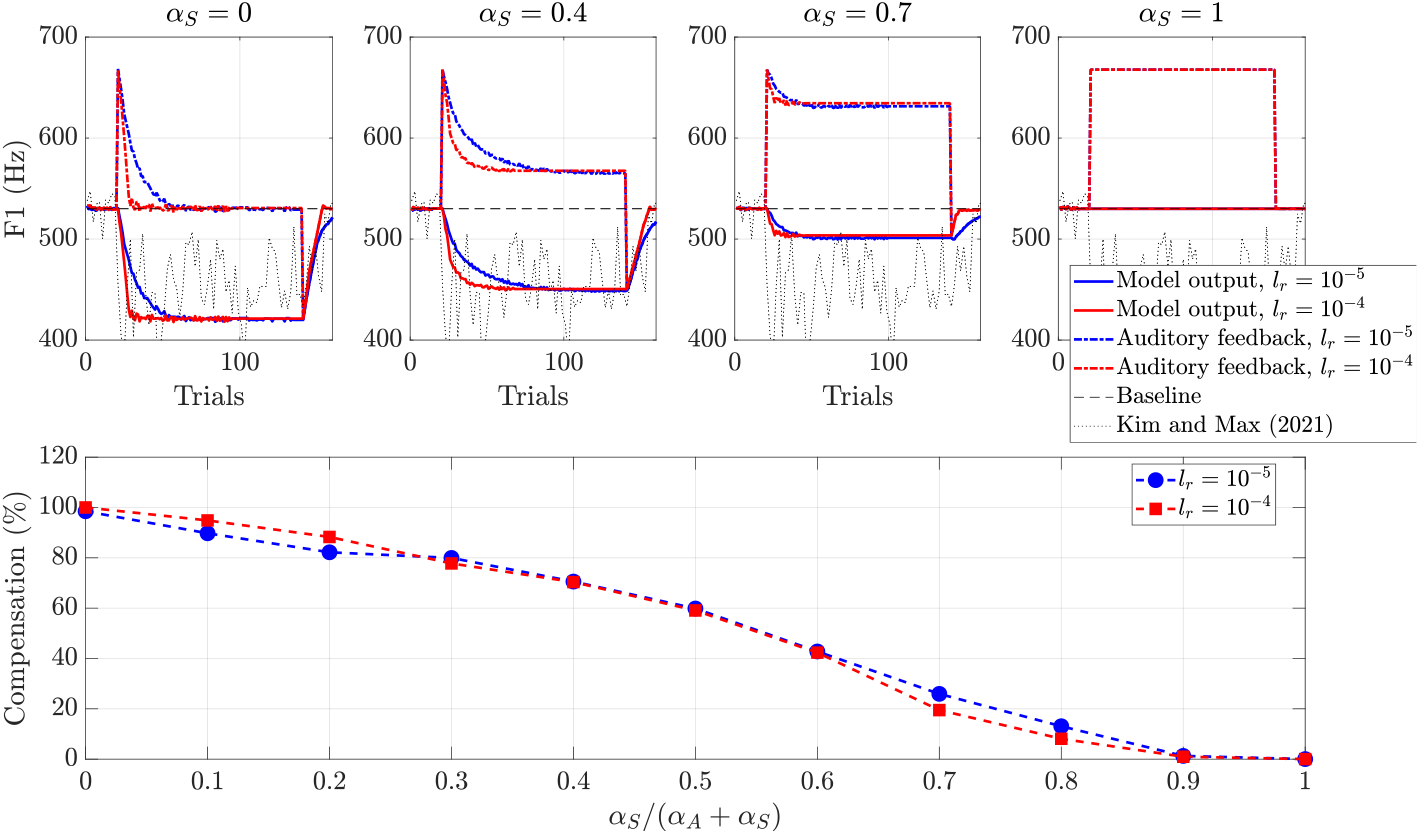
Top panel: The evolution of produced first formant frequency (solid lines) during simulated trials for different values of *α*_*S*_, the weight assigned to the somatosensory intelligibility and 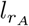, the learning rate of the auditory internal model. From left to right: *α*_*S*_ = 0 (only acoustic intelligibility is taken into account), *α*_*S*_ = 0.4, *α*_*S*_ = 0.7, and *α*_*S*_ = 1 (only somatosensory intelligibility is taken into account). The dash-dotted lines represents auditory feedback, and the dashed black line represents the baseline value of F1. The dotted black line corresponds to an example of experimentally recorded adaptation behavior from Kim and Max [21]. The bottom panel shows the compensation ratio, expressed in %, returned by the simulations as a function of *α*_*S*_ for two values of the learning rate, 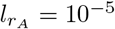 and 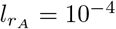.

Our simulations show that the balance between acoustic and somatosensory intelligibility optimization has an effect on the level of adaptation to altered auditory feedback: the greater the priority assigned to acoustic intelligibility, the larger the compensation. With maximal priority assigned to acoustic intelligibility (*α*_*S*_ = 0, *α*_*A*_ = 1), our model predicts full compensation, as the perceived first formant frequency during the auditory feedback altered trials is equal to the first formant frequency during the non-perturbed trials. With increasing *α*_*S*_, the compensation degree gradually decreases. Finally, when *α*_*S*_ = 1 (and *α*_*S*_ = 0, *i*.*e*. acoustic intelligibility is not optimized), there is no adaptation at all.

Figure 4 also shows simulated adaptation behavior for two different values of the learning rate 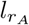 of the auditory internal model, *i*.*e*. the amount of correction to apply to the internal auditory forward model’s weights during the update at each epoch. It shows that the learning rate controls the speed at which the speaker reaches its maximal adaptation. For a higher learning rate (here 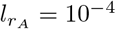), it takes fewer trials for the model output to reach maximal compensation than for a lower learning rate (here 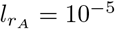). There is a small impact of learning rate on compensation, but this depends systematically on the relative weighting of acoustic and somatosensory intelligibility. For *α*_*S*_ *<* 0.5, a higher learning rate yields slightly more compensation, while for *α*_*S*_ *>* 0.5, a lower learning rate yields slightly more compensation. Combination of the dual requirement of predicting and continuous adapting of the internal model of both acoustic and somatosensory consequences of articulatory action thus answers Objective B of incomplete compensation as well as potentially the Objective C of inter-speaker variability. This is because it is possible that values of the weights *α*_*A*_ and *α*_*S*_ are speaker-dependent. That is, speakers adapt differently to altered sensory feedback because they have different values of *α*_*A*_ and *α*_*S*_. Similarly, speakers adapt more or less quickly to altered sensory feedback because their internal forward models have different learning rates.

### The influence of the weights assigned to the effort cost

Figure 5 shows the results of our simulations for the American-English vowel /ε/ under formant-shifted perturbations shown in Figure 1, for different values of *α*_*E*_, the weight assigned to the articulatory effort cost. This parameter is directly related to the level of planned hyper- and hypo-articulation [7, 8]: the lower the weight assigned to the effort cost, the greater the predicted hyperarticulation.

**Figure 5.**
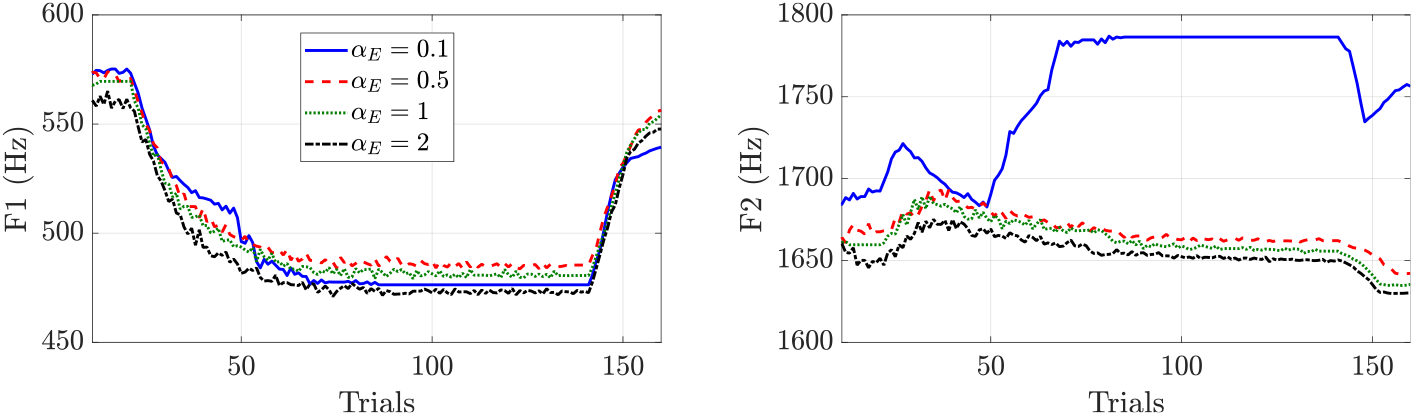
Evolution of the simulated produced formant frequencies as a function of trials for different values of *α*_*E*_, at fixed values of *α*_*A*_ = 0.6 and *α*_*S*_ = 0.4. The left panel shows F1 and the right panel shows F2.

Our simulations predict that the degree of planned hyper-articulation slightly changes the adaptation response of the speaker. This is evidenced by the left panel of Figure 5, which shows produced F1 for different values of *α*_*E*_. Note that increasing *α*_*E*_ yields a lower F1 baseline (measured as the produced F1 averaged across trials 11 to 20. This is because the baseline solution with high *α*_*E*_ yields a more centralized /ε/, namely it is closer to the neutral solution **x** = **0**, which generates *F* 1 = 417 Hz. The compensation ratio computed from the simulations shown in Figure 5 are 80%, 71%, 73%, and 72%, for *α*_*E*_ = 0.1, *α*_*E*_ = 0.5, *α*_*E*_ = 1, and *α*_*E*_ = 2, respectively. These different compensation ratios indicate that, although the degree of planned hyper-articulation changes the degree of compensation, there is no direct rank correlation between these two variables. The right panel of Figure 5 shows the effect on F2. It shows that hyperarticulation (*i*.*e*. low *α*_*E*_) favors changes in F2: *α*_*E*_ = 0.1 yields a rise of F2 of about 5%, while it is slightly raised before going down towards the baseline for higher values of *α*_*E*_. This influence of the degree of hypo- and hyper-articulation on unaltered formants during adaptation, as predicted by our approach, might explain variability which has been observed in previous experiments. Indeed, some studies reported modifications of F2 during F1-altered feedback experiments [11, 14, 16, 29], while another study did not report significant changes [55]. This could be due to inter-speaker variability of hyper-articulation across these different experiments. In that case, our approach can also simulate inter-speaker (or intra-speaker) variability of adaptation behavior (which is our Objective C) by changing the weight assigned to the effort cost, *i*.*e*. by varying the degree of planned hyper-articulation.

### The influence of the architecture and training of the internal models

In our approach, speech adaptation behavior to altered auditory feedback depends on the ability of the internal models to learn a new relationship between motor commands and acoustics during the auditory feedback perturbation phase. This can be seen in Figure 4, which shows the effect of the learning rate of the auditory internal model on adaptation behavior. Another possible variable that may impact adaptation behavior is the amount of data on which internal models have been trained, and also the general architecture (number and size of hidden layers). In order to investigate this potential effect, we ran simulations of adaptation experiments using various pre-trained internal forward models *𝒜*^(0)^ and *𝒮*^(0)^, each differing according to the size of the dataset used to train them, and also on the number and size of hidden layers.

Figure 6 shows the results of these experiments for these different pre-trained internal models, with fixed *α*_*E*_ = 0.5, *α*_*S*_ = 0.4 and *α*_*A*_ = 0.6, and fixed learning rate of 10^*−*5^. Results show that the architecture of the internal models and how internal models have been trained have an influence on adaptation behavior. For instance, for fixed effort and auditory weights, and fixed learning rates, the number of training samples affects the amount of compensation (speakers compensate more with models trained on more data), in addition to the adaptation rate (the speaker reaches their maximal compensation faster with internal models trained on less data). In other words, our approach predicts that speakers with less experience in speaking compensate less, but faster, than speakers with more speaking experience. However, similarly to the effect of the degree of planned hyper-articulation, there is no direct rank correlation between size of training data and amount of compensation, as the compensation ratio obtained with models pre-trained on 1e4 samples is larger than the compensation ratio obtained with models pre-trained on more data (5e4 and 1e5 samples). Regarding the architecture of the internal models, larger internal models (more hidden layers and larger hidden layers) yield faster adaptation behavior with higher compensation amplitude than smaller internal models.

**Figure 6.**
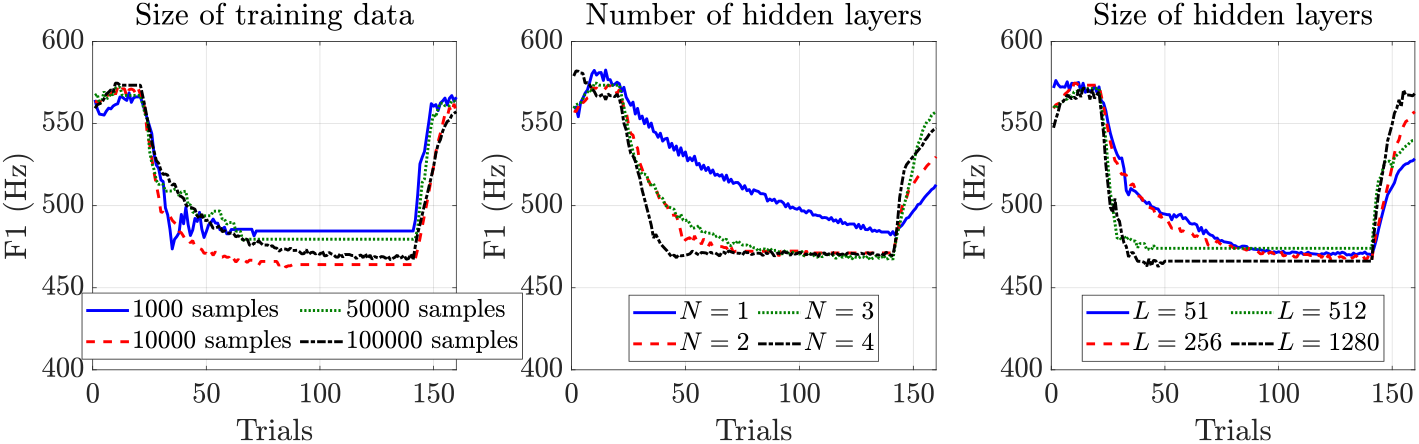
The evolution of the simulated produced first formant frequency F1 as a function of trial number for different internal models. The left panel shows the effect of the number of samples used for training internal models with nominal architectures (100,000 samples are the nominal models, as presented in Table 1). The central panel shows the effect of the number of hidden layers (*N* = 3 are nominal models) and the right panel shows the effect of the size of hidden layers (*L* = 256 are nominal models).

These results show that different characteristics of internal forward models can be a source of inter-speaker variability and, consequently, these observations answer Objective C. However, the characteristics of internal forward models only impact the compensation ratio and the compensation rate of speakers, but not the general compensation patterns. These different pre-trained internal forward models did not generate unusual adaptation behavior, such as no compensation or compensation in a different direction (For example, F1 that goes in the same direction as the perturbation). In addition, it is worth noting that the compensation ratio and the compensation rate can be adjusted or modified by other means, such as by the degree of planned hyper-articulation (defined by the ratio between weights assigned to effort and recognition costs), the learning rate, or by adjusting the relative contributions of auditory and somatosensory recognition costs in the global cost function.

## Results of simulations with altered somatosensory feedback

Modeling adaptation to altered somatosensory feedback using our approach is relatively similar. The major difference is that perturbation of articulatory movements, e.g., perturbation of jaw movements, as proposed in [13], also impacts produced acoustics. As a consequence, both auditory and somatosensory feedback received by the speaker differ from predicted sensory feedback. However, since our approach does not require the detection or the evaluation of discrepancies between predicted and actual sensory feedback, our model deals with altered somatosensory feedback in exactly the same way as with altered auditory feedback.

In this section, we present simulations which are inspired by Lametti *et al*. [13]. The aim of this section is to show that, accordingly to our Objective E, our approach is also able to reproduce adaptation to altered somatosensory feedback. We model somatosensory perturbation by artificially pulling the jaw outward, horizontally, and by artificially increasing the lip protrusion. This is done by modifying the *x*-coordinates of the upper and lower contours of the vocal tract given by the Maeda model from the jaw to the end of the lips: the horizontal distance *d* of the jaw from a reference point (here the center of the polar grid used to compute the area function, located at *x* = 10) is multiplied by the perturbation factor *k*, and lip protrusion is also multiplied by *k*. Figure 7 shows an example of such geometry perturbation for *k* = 1.26. The series of perturbations is similar to the one used for the altered auditory feedback simulations, namely as shown in the right panel of Figure 7.

**Figure 7.**
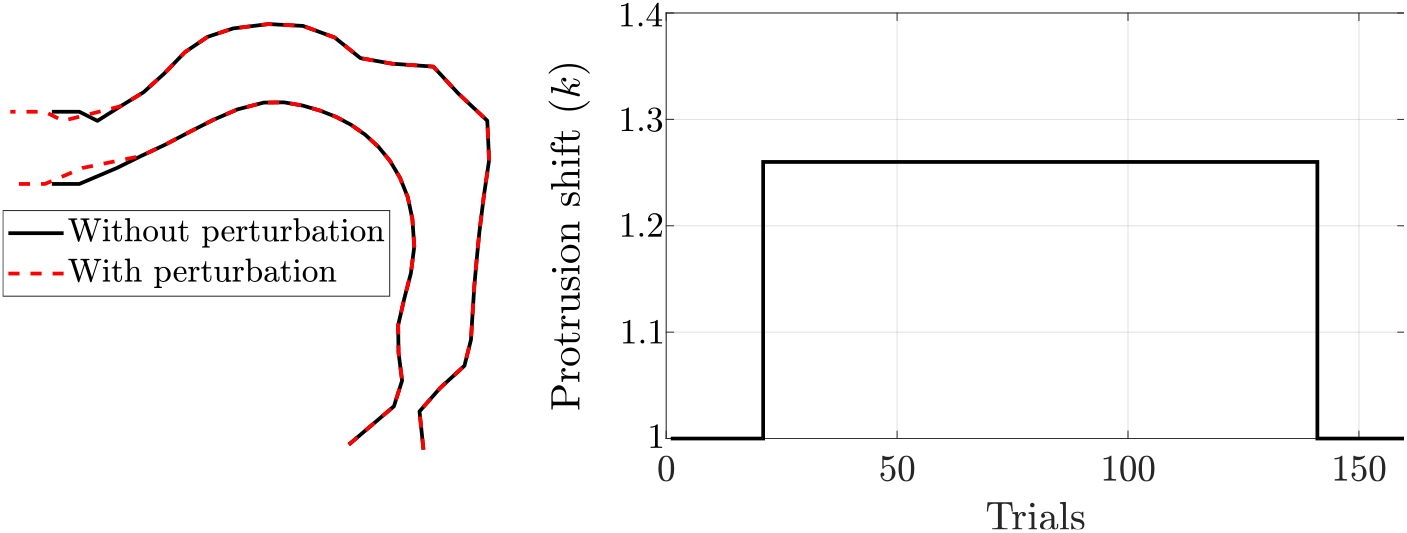
Example of jaw perturbation with *k* = 1.26 (left panel) and perturbation shift *k* as a function of trials used for the experiment (right panel).

Figure 8 shows the results of our simulations for three values of the weights assigned to auditory and somatosensory recognition costs. In the purely-auditory planning case (*α*_*S*_ = 0, *α*_*A*_ = 1), represented in the left panels in Fig. 8, the change in F1 induced by the articulatory perturbation is fully compensated: the lowering of F1 induced by the jaw pulling (which lengthens the vocal tract, hence reduce F1) is compensated as the produced F1 goes back to the value before perturbation around trial 60. The lip protrusion perturbation is, however, not fully compensated, as shown in the bottom left plot in Fig. 8: at trial 60, when the produced F1 is back to the value before perturbation, the optimal chosen motor command is such that the produced lip protrusion is still higher than before perturbation (around 1.1 cm, while it is 1 cm before perturbation). After the perturbation is applied, namely after trial 140, we also observe an acoustic adaptation. In the absence of articulatory perturbation, the produced F1 is now too high, but the produced F1 goes back towards the baseline in subsequent trials (not quickly enough to reach full compensation before the end of the simulated experiments). Similarly, the chosen optimal motor commands are such that the lip protrusion is increased such that lip protrusion tends towards the value of 1 cm, namely that before the first perturbations. In the purely somatosensory-based planning case (*α*_*S*_ = 1, *α*_*A*_ = 0), which is represented in the right panels in Fig. 8, the change in F1 induced by the articulatory perturbation is not fully compensated. However, lip protrusion is fully compensated: during the adaptation phase, the optimal motor commands are chosen such that the perturbed lip protrusion goes back to the baseline value. In the intermediate case (*α*_*S*_ = 0.5, *α*_*A*_ = 0.5), as shown in the central panels in Fig. 8, our model predicts incomplete compensation for both F1 and lip protrusion.

**Figure 8.**
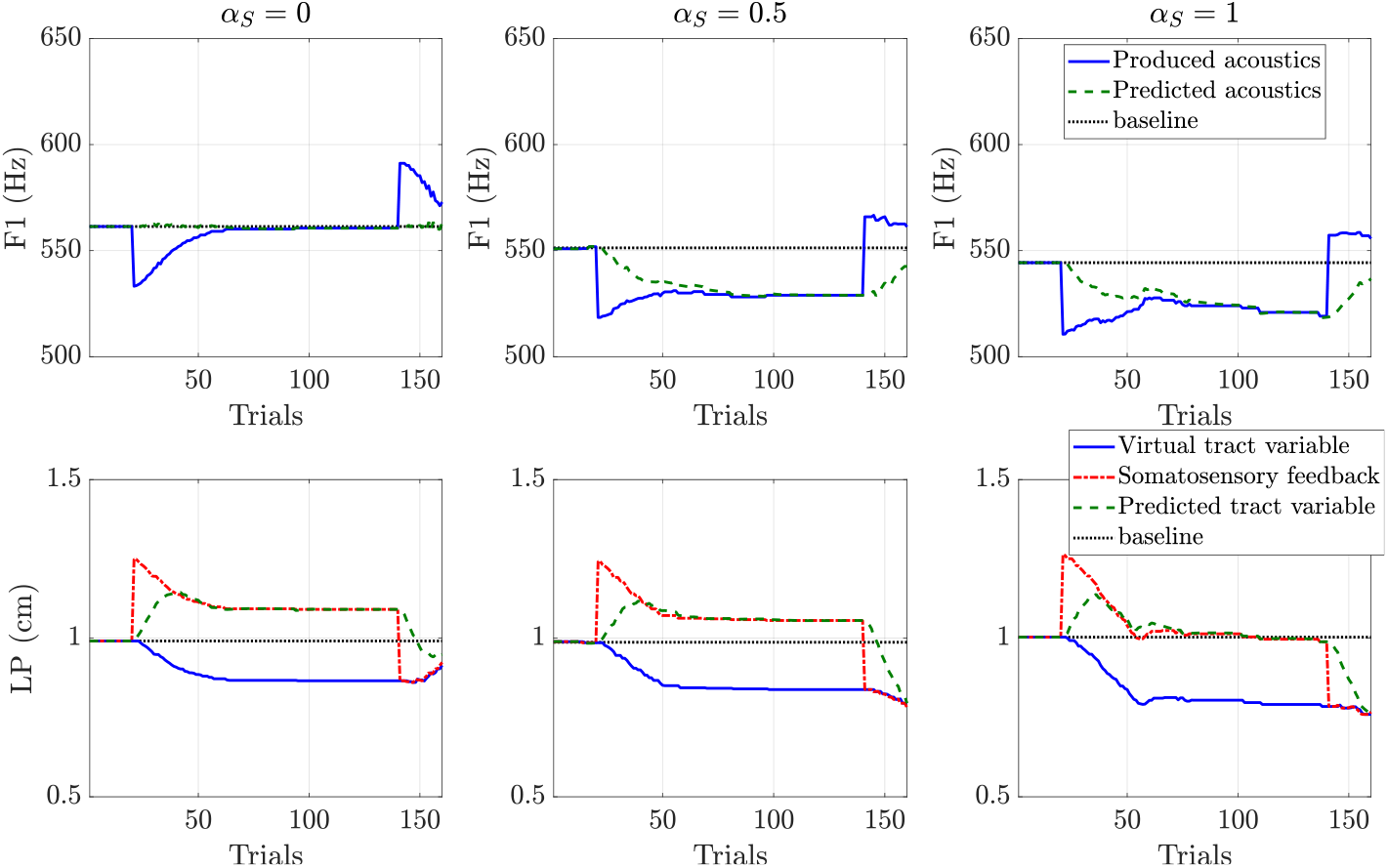
Simulated adaptation to altered somatosensory feedback for three values of *α*_*S*_, namely *α*_*S*_ = 0 (left panels), *α*_*S*_ = 0.5 (central panels), and *α*_*S*_ = 1 (right panels). The top panels show simulated acoustic output (first formant frequency F1), including produced acoustics (blue solid line) and acoustics predicted by the auditory internal model (dashed green line). The bottom panels show simulated tract variable lip protrusion (LP), including virtual lip protrusion (without perturbation, solid blue line), received somatosensory feedback (after perturbation, dotted-dashed red line), and lip perturbation predicted by the somatosensory internal model (dashed green line).

### Fit to experimental data

The previous section showed that our approach was able to reproduce several observed aspects of adaption to altered auditory feedback qualitatively, including gradual and incomplete compensation, and inter- and intra-speaker variability. This section investigates the ability of our approach to reproduce these behaviors quantitatively. In other words, how well can the acoustic output generated by our model fit observed experimental data? For that purpose, for each of the F1 trajectories recorded from the 24 speakers of the Kim and Max experiments [21], we estimated some parameters of our model for which the simulated F1 output best fits the experimental F1 trajectories. We chose to estimate only two parameters, namely the auditory weight *α*_*A*_ and the learning rate of the internal auditory model 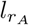, because our simulations with different weights *α*_*E*_ showed that this parameter had a small impact on characteristics (compensation ratio and compensation rate, as shown in Fig. 5) that can also be adjusted via modifications of *α*_*A*_ and 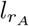, as shown in Fig. 4.

The fit is done by minimizing an objective function which is the Normalized Mean Root Square Error (NRMSE) between the fit (the F1 trajectory generated by the model) and the experimental data (F1 produced by the speaker, as reported in [21]), where the fit and the experimental data are each normalized by their baseline values (taken as the mean between trials 11 and 20). For each speaker, the minimization is done running 100 instances of the Nelder-Mead algorithm [65] with random initialization of *α*_*A*_ and 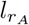. For each speaker, the final estimate of *α*_*A*_ and 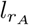 is the solution of the Nelder-Mead minimization which returns the lowest NRMSE across the 100 runs. In addition, we compare the results of our fit with the fit obtained with SimpleDIVA [39], because it is the only adaptation model which has been used to fit and analyze experimental data.

### A 2-parameter version of SimpleDIVA

The SimpleDIVA model, as presented in [39], is a simple 3-parameter model based on the DIVA model [40–42]. It is used to model feedback and feedforward control mechanisms involved in the production of formant trajectories. It has been developed to model and fit data from adaptation to altered auditory feedback experiments. The model simply assumes that the formant pattern produced by the speaker at a given trial *n* is the sum of the feedforward command and a sensory feedback-based correction, hence:

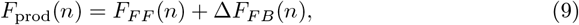

where *F*_prod_(*n*) is the produced formant at trial *n, F*_*F F*_ (*n*) is the feedforward command, and Δ*F*_*F B*_(*n*) is the sensory feedback-based correction. The feedback-based correction is a weighted sum of the contributions of both auditory and somatosensory feedback, as:

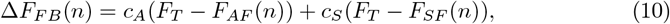

where *c*_*A*_ and *c*_*S*_ are coefficients that specify the contributions of auditory and somatosensory feedback, respectively, in the feedback-based correction, *F*_*T*_ is the vector containing the target formants, and *F*_*AF*_ (*n*), and *F*_*SF*_ (*n*) are formants corresponding to the auditory and somatosensory feedback received at trial *n*, respectively. SimpleDIVA assumes that the feedforward command *F*_*F F*_ is regularly updated based on the feedback received on the preceding trial. The equation for updating *F*_*F F*_ is

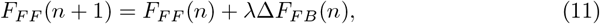

where *λ* is a parameter which controls the rate at which adaptation is performed by the speaker.

The three parameters of the SimpleDIVA model are then *c*_*A*_, *c*_*S*_, and *λ*. Optimization procedures can be used to fit the model to real experimental data [39]. These consist of finding values for the three parameters of the SimpleDIVA model for which the generated formant trajectories best fit the formant trajectories extracted from the experimental recordings. This has been done, for instance, to compare the values of these parameters between different populations, such as adults and children, and children with complex speech sound disorders [24].

From Eqs. (10) and (11), one can see that the *λ* parameter multiplies the coefficients *c*_*A*_ and *c*_*S*_ by a common factor, which makes the *λ* parameter mathematically redundant. That is, the SimpleDIVA model can be reduced to a simple 2-parameter model by setting *λ* = 1. Doing so reduces the complexity of the fit and ensures the uniqueness of the solution.

## Results of the fits

Table 2 presents the results of the fits for both our approach (XT/3C) and SimpleDIVA [39], and Figure 9 shows the comparison between the generated F1 trajectories obtained from the fit with our approach and SimpleDIVA, and the experimental F1 trajectories from [21]. The fit error (column NRMSE) shows very similar values for both XT/3C (mean NRMSE is 0.0442) and SimpleDIVA (mean NRMSE is 0.0436), as well as similar Pearson correlation coefficient (mean *r* is 0.40 for XT/3C, 0.43 for SimpleDIVA) for all speakers. This similar ability of XT/3C and SimpleDIVA to fit experimental trajectories is illustrated in Figure 9, as the F1 trajectories generated by both models are very close to each other. This shows that our approach can quantitatively reproduce adaptation behavior with performance that is comparable to that of SimpleDIVA.

**Table 2.** Results of fits per speakers and method.

| Speakers | XT/3C |  |  |  | SimpleDIVA |  |  |  |
| --- | --- | --- | --- | --- | --- | --- | --- | --- |
| | $\alpha_A$ | $l_{r_a}$ | NRMSE | $r$ | $c_A$ | $c_S$ | NRMSE | $r$ |
| S01 | 0.84 | 1.2e-05 | 6.06e-02 | 0.26 | 5.78e-02 | 2.16e-02 | 5.90e-02 | 0.28 |
| S02 | 0.77 | 7.6e-04 | 5.77e-02 | 0.52 | 1.72e-01 | 9.57e-02 | 5.25e-02 | 0.61 |
| S03 | 0.50 | 5.5e-05 | 4.19e-02 | 0.65 | 1.32e-01 | 1.82e-01 | 3.84e-02 | 0.72 |
| S04 | 0.50 | 1.0e-05 | 3.75e-02 | 0.55 | 1.04e-02 | 1.42e-02 | 3.63e-02 | 0.58 |
| S05 | 0.51 | 1.0e-03 | 3.70e-02 | 0.15 | 2.80e-03 | 2.70e-01 | 3.73e-02 | 0.13 |
| S06 | 0.58 | 7.1e-05 | 3.56e-02 | 0.76 | 3.57e-02 | 3.37e-02 | 3.68e-02 | 0.77 |
| S07 | 0.56 | 9.1e-05 | 3.13e-02 | 0.68 | 5.51e-02 | 3.39e-02 | 2.97e-02 | 0.69 |
| S08 | 0.57 | 7.4e-05 | 4.75e-02 | 0.51 | 5.68e-02 | 3.56e-02 | 4.70e-02 | 0.52 |
| S09 | 0.62 | 1.0e-03 | 3.45e-02 | 0.30 | 1.61e-02 | 5.79e-02 | 3.48e-02 | 0.27 |
| S10 | 0.71 | 1.0e-05 | 3.53e-02 | 0.62 | 3.88e-02 | 2.02e-02 | 3.49e-02 | 0.62 |
| S11 | 0.53 | 1.0e-05 | 4.71e-02 | 0.39 | 2.34e-02 | 3.03e-02 | 4.70e-02 | 0.39 |
| S12 | 0.46 | 4.3e-05 | 4.58e-02 | 0.44 | 3.72e-02 | 1.06e-01 | 4.60e-02 | 0.44 |
| S13 | 0.15 | 1.3e-05 | 3.60e-02 | 0.14 | 3.33e-03 | 2.95e-02 | 3.48e-02 | 0.33 |
| S14 | 0.48 | 9.1e-04 | 5.16e-02 | 0.29 | 1.09e-01 | 1.45e+00 | 5.13e-02 | 0.31 |
| S15 | 0.15 | 1.1e-05 | 6.41e-02 | 0.22 | 7.76e-04 | 3.23e-03 | 6.40e-02 | 0.14 |
| S16 | 0.79 | 9.7e-04 | 4.78e-02 | 0.38 | 2.25e-02 | 3.33e-02 | 4.85e-02 | 0.35 |
| S17 | 0.33 | 1.0e-05 | 3.99e-02 | 0.04 | 1.33e-03 | -6.04e-03 | 3.77e-02 | 0.33 |
| S18 | 0.55 | 2.5e-04 | 4.29e-02 | 0.46 | 3.55e-02 | 5.04e-02 | 4.28e-02 | 0.46 |
| S20 | 0.40 | 8.8e-05 | 3.68e-02 | 0.32 | 1.49e-02 | 6.40e-02 | 3.67e-02 | 0.32 |
| S21 | 0.37 | 5.2e-04 | 4.61e-02 | 0.23 | 9.01e-03 | 7.75e-02 | 4.62e-02 | 0.24 |
| S22 | 0.80 | 4.5e-04 | 8.08e-02 | 0.23 | 2.63e-02 | 1.93e-02 | 8.10e-02 | 0.24 |
| S23 | 0.39 | 5.9e-05 | 2.66e-02 | 0.57 | 2.57e-02 | 1.54e-01 | 2.68e-02 | 0.57 |
| S24 | 0.64 | 9.9e-04 | 3.27e-02 | 0.47 | 1.44e-02 | 4.25e-02 | 3.23e-02 | 0.49 |
| Mean | 0.53 | 3.2e-04 | 4.42e-02 | 0.40 | 3.75e-02 | 1.17e-01 | 4.36e-02 | 0.43 |

**Figure 9.**
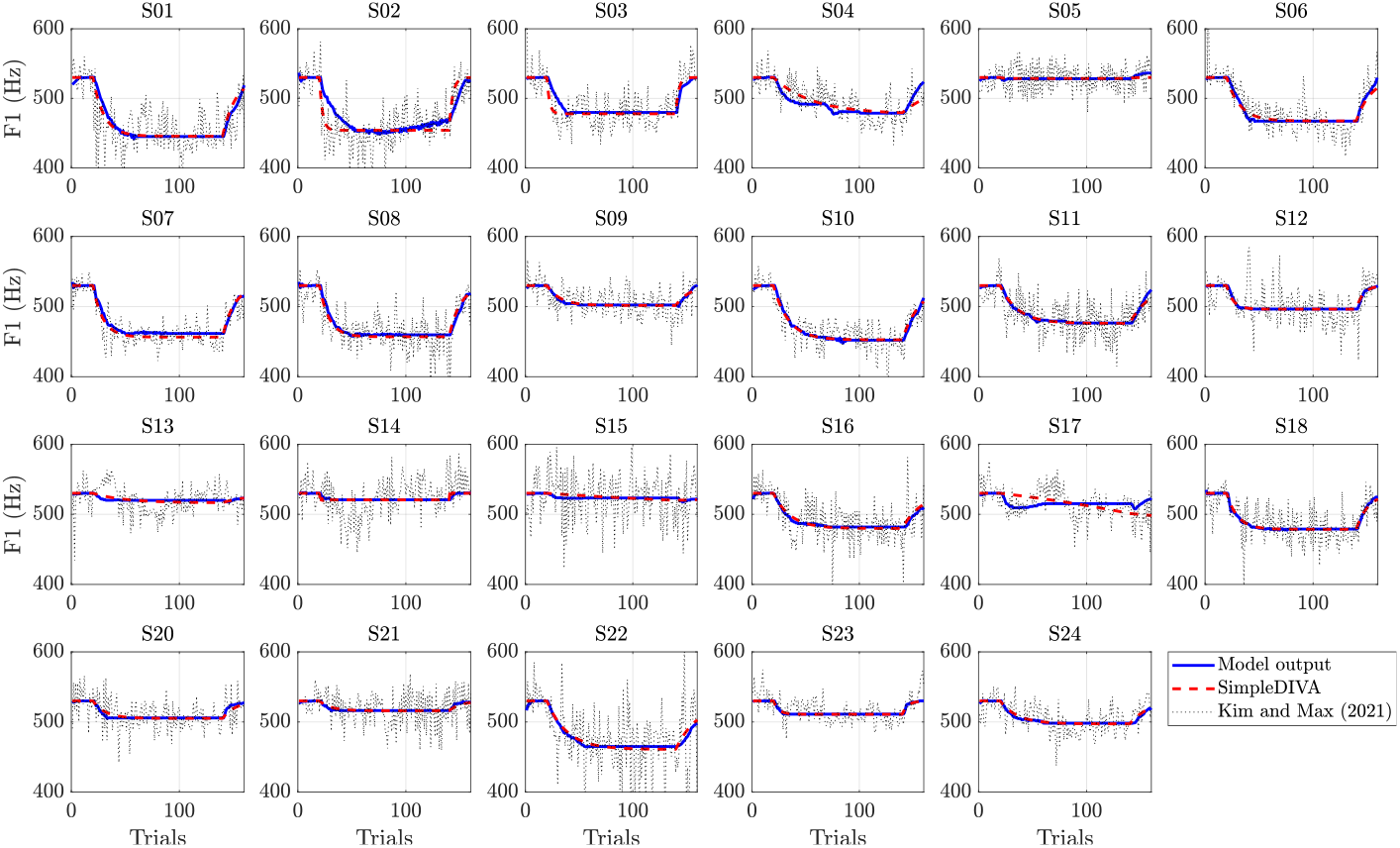
Results of best fits obtained with SimpleDIVA (red dashed line) and XT/3C (blue solid line) for the 24 speakers.

Interestingly, we found strong correlations between the estimated parameters of our model and the estimated parameters of SimpleDIVA, as shown in Figure 10. The weight *α*_*A*_ assigned to the auditory recognition cost in our approach (relatively to *α*_*A*_ + *α*_*S*_) and *c*_*A*_ the auditory gain in of SimpleDIVA (relatively to *c*_*A*_ + *c*_*S*_) are correlated with a Pearson correlation coefficient of 0.65 (*p <* 0.001). The somatosensory weight *α*_*S*_ in our approach and the somatosensory gain *c*_*S*_ of SimpleDIVA are similarly correlated. This is interesting given that despite some obvious architectural similarities, the two models are based on different principles. The key difference being that SimpleDIVA updates the feedforward command by a compensation factor proportional to the difference between the feedback and the target, while our approach updates the feedforward command because the estimation of the optimal solution of the cost function changes due to the update of the speaker’s internal forward models.

**Figure 10.**
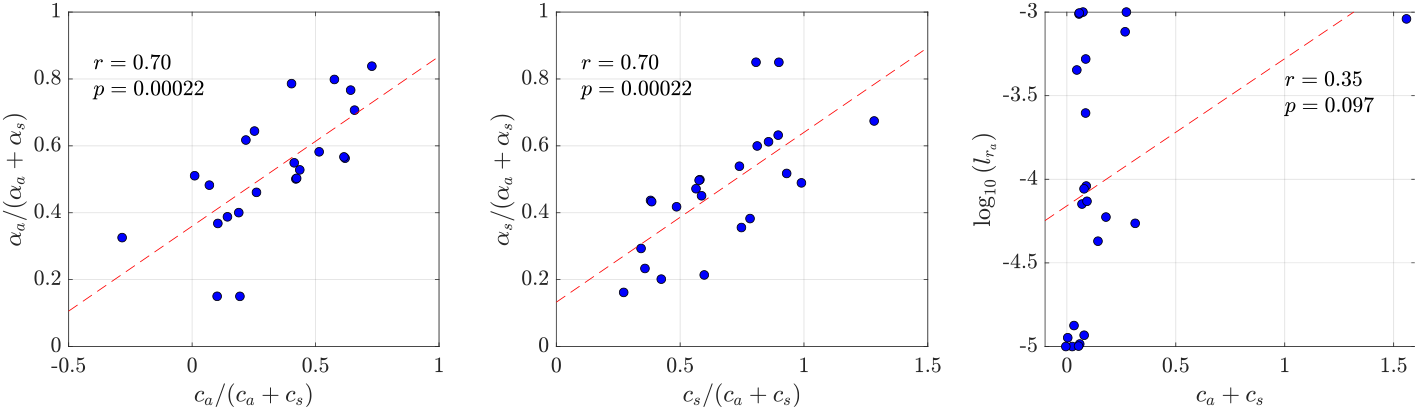
Estimated parameters of our model (*α*_*E*_, *α*_*A*_ and 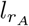) as a function of the estimated SimpleDIVA parameters *c*_*A*_ and *c*_*S*_ for the 24 experimental F1 trajectories taken from [21].

## Conclusion

This paper has presented a development of the optimization-based model of speech production, derived from [6–9], that forms the Phonetic Planning component of Turk and Shattuck-Hufnagel’s XT/3C model [1, 2]. This new version is an extension of the original model, where extensions include adding a somatosensory-based loop of intelligibility prediction and a continuous update of internal sensory predictions based on sensory feedback. We have shown that our new model accounts for the key acoustic and articulatory features of motorsensory adaptation of speech to altered sensory feedback. More precisely, these key features include gradual (Objective A) and incomplete adaption (objective B), inter-speaker variability (Objective C), adaptive modification in unaltered formants (Objective D), and articulatory adaptation to altered somatosensory feedback (Objective E).

In our approach, the mechanism of motorsensory adaptation is based on regular and continuous updates of the speaker’s internal models that drive the Motor-Sensory Implementation stage of speech production, based on sensory feedback perceived by the speaker. The updated internal models are used during the optimization of the cost function that occurs during Phonetic Planning for any subsequent production. The internal models that require updates to account for compensation and adaptation behavior are 1) a model that predicts the acoustic consequences of motor (articulatory commands) and 2) a model that predicts the somatosensory sensations (here tract variables) from given motor commands. The idea of continuous updates of internal models is compatible with the suggestions of [66] and provides appropriate mechanisms for flexible and continuous learning of speech production. Internal models of the relationship between articulation and their sensory consequences are proposed to develop from infancy; flexible and continuous updates of these models are required to account for adjustments to speech articulation in reaction to changes in vocal tract anatomy (during growth, after surgery, or during temporary speech disorders), as well as in reaction to a changing external environment (e.g. environmental noise, or altered feedback in experimental situations).

We have modeled these internal forward models as Artificial Neural networks (ANN). The weights of these ANN internal forward models are updated at each iteration, using the motor commands passed to the plant and the feedback that result from these motor commands. Phonetic planning (cost-function minimization) that makes use of these internal models successfully reproduces gradual motor-sensory adaptation to altered feedback, both when auditory feedback is altered (F1 shifted), and when somatosensory feedback is altered (forced jaw and lip protrusions).

When only the auditory feedback was perturbed, the optimization-based model of speech production returned articulatory output for which the first formant frequency moved gradually in the direction opposite to the perturbation, fulfilling Objective A. When jaw and lip protrusion were perturbed (artificially increased), the model predicted a compensation in the motor commands that resulted in perturbed jaw and lip protrusion at the end of the perturbation phase which were closer to the baseline than at the beginning of the perturbation phase. This shows that adaptation to altered somatosensory feedback could also be modeled using our approach, as required by Objective E. Our simulations also show that the rate at which the adaptation was made could be controlled by adjusting the learning rate of the internal models. Basically, the higher the learning rate, the faster the adaptation. Regarding inter-speaker variability, which is Objective C, this shows that different adaptation behavior can be controlled and modeled with our approach via modifications of meaningful model’s parameters.

The simulation results with and without a somatosensory intelligibility criterion showed the necessity of having both auditory and somatosensory intelligibility costs in our model. More specifically, using a single auditory-based intelligibility criterion incorrectly results in complete (or almost complete) adaptation, whereas, observations in the literature consistently show incomplete adaptation (e.g., in [12, 19, 21, 23, 54]). We therefore introduced an intelligibility cost based on somatosensory characteristics, to be used alongside the intelligibility cost based on acoustic (or auditory) characteristics, which we had used in our previous studies [6–9]. This extension was inspired by feedback-based models of speech production, which generally use two sensory feedback subsystems, to account for auditory and somatosensory feedback [36, 37, 39, 41–43, 45], and is conceptually most similar to Bayesian GEPPETO [36, 37, 43].

Our simulations showed that the introduction of this additional intelligibility score in the cost function successfully reproduced the phenomenon of incomplete compensation, which fulfills Objective B. The compensation rate could be modified by adjusting the relative weights assigned to both intelligibility functions, which provides another source of inter-speaker variability. Additionally, our simulations highlighted another source of variability, namely the ratio between the weights assigned to effort vs. recognition costs, which result in different degrees of hyper-vs. hypo-articulation. Indeed, our model predicts that modifying the weight assigned to the effort cost, which is directly related to patterns of hyper-/hypo-articulation, changes adaptation behavior. To the best of our knowledge, this is the first study of the impact of planned hyper-articulation on adaptation behavior. Consequently, this observation requires further experimental investigation for validation. These results show that our model can successfully simulate inter-speaker variability of adaptation behavior, as required by Objective C. This is because the characteristics of this adaptation behavior can be modified by adjusting the parameters of our model: the learning rate of the internal models impacts adaptation speed and the weights assigned to the auditory and somatosensory intelligibility tasks in the cost function impact the amount of compensation during adaptation.

Finally, we presented an evaluation of the approach in terms of model fits to experimentally observed adaptation behavior to altered auditory feedback. This consisted of estimating the model parameters for which our model output fit best experimental data taken from [21]. The results of this evaluation shows that our approach can accurately fit real F1 trajectories recorded during adaptation experiments. The fit error is very similar to the fit error obtained with SimpleDIVA [39], the only adaptation model which has been used to fit and analyze experimental data. Our results also showed that the parameters of our model were correlated with the parameters of SimpleDIVA.

The weights assigned to auditory and somatosensory intelligibility tasks are correlated with the gains of the auditory and somatosensory feedback control subsystems, respectively. Similarly to SimpleDIVA [39], this inversion method (estimating the input parameters of a forward model from observations of its outputs) presents the interest of interpreting the results of adaptation experiments in terms of speakers’ sensory preference and learning rate, that is with a few number of meaningful and interpretable parameters. In addition, unlike SimpleDIVA, which only predicts acoustic consequences of altered acoustic feedback during adaptation experiments, our approach offers the possibility of investigating the articulatory consequences of altered motorsensory feedback. It would be interesting in the near future to compare our predictions of articulatory compensation with real data taken from perturbation experiments.

## Acknowledgments

This research has received funding from the European Research Council (ERC) under the European Union’s Horizon 2020 research and innovation programme (Grant agreement No. 101019847).

## Footnotes

1 Note that we could also include a duration cost to account for the brevity requirement, as proposed in [5, 6, 48–50]. For the sake of simplicity, this is not included in this paper, mainly because we will simulate static production of vowels

2 Note that we recently showed in [61] that our model can potentially accept intelligibility based on a wider stretch of speech.

3 The experimental data have been extracted from the authors’ repository at https://osf.io/s84df/.

